# Epithelial retinoic acid receptor β regulates serum amyloid A expression and vitamin A-dependent intestinal immunity

**DOI:** 10.1101/563908

**Authors:** Sureka Gattu, Ye-Ji Bang, Mihir Pendse, Chaitanya Dende, Andrew L. Chara, Tamia A. Harris, Yuhao Wang, Kelly A. Ruhn, Zheng Kuang, Shanthini Sockanathan, Lora V. Hooper

## Abstract

Vitamin A is a dietary component that is essential for the development of intestinal immunity. Vitamin A is absorbed and converted to its bioactive derivatives retinol and retinoic acid by the intestinal epithelium, yet little is known about how epithelial cells regulate vitamin A-dependent intestinal immunity. Here we show that epithelial cell expression of the transcription factor retinoic acid receptor β (RARβ) is essential for vitamin A-dependent intestinal immunity. Epithelial RARβ activated vitamin A-dependent expression of serum amyloid A (SAA) proteins by binding directly to *Saa* promoters. In accordance with the known role of SAAs in regulating Th17 cell effector function, epithelial RARβ promoted IL-17 production by intestinal Th17 cells. More broadly, epithelial RARβ was required for the development of key vitamin A-dependent adaptive immune responses, including CD4^+^ T cell homing to the intestine and the development of immunoglobulin A-producing intestinal B cells. Our findings provide insight into how the intestinal epithelium senses dietary vitamin A status to regulate adaptive immunity and highlight the role of epithelial cells in regulating intestinal immunity in response to diet.

**Significance Statement:** Vitamin A is a nutrient that is essential for the development of intestinal immunity. It is absorbed by gut epithelial cells which convert it to retinol and retinoic acid. Here we show that the transcription factor retinoic acid receptor β (RARβ) allows epithelial cells to sense vitamin A in the diet and regulate vitamin A-dependent immunity in the intestine. We find that epithelial RARβ regulates several intestinal immune responses, including production of the immunomodulatory protein serum amyloid A, T cell homing to the intestine, and B cell production of immunoglobulin A. Our findings provide new insight into how epithelial cells sense vitamin A to regulate intestinal immunity and highlight why vitamin A is so important for immunity to infection.

---

The mammalian intestinal epithelium is a vital interface between the external environment and internal tissues. Epithelial cells interact with the environment of the gut lumen by absorbing dietary compounds and by associating with the resident bacterial communities that promote digestion. The intestinal epithelium also orchestrates development of the underlying immune system through the secretion of immunoregulatory proteins (1). Thus, epithelial cells are ideally positioned to capture information about the diet and the microbiota in order to regulate adaptive immunity. While it is known that gut epithelial cells detect intestinal microorganisms through various pathways involving pattern recognition receptors (1), little is known about how epithelial cells sense dietary components to regulate adaptive immunity.

Vitamin A is a fat-soluble nutrient that is essential for the development of adaptive immunity to intestinal microorganisms. It is required for immunoglobulin A (IgA) production by intestinal B cells (2), T cell homing to the intestine (3), and the production of interleukin-17 (IL-17) by T helper 17 (Th17) cells (4). Consequently, vitamin A-deficient diets result in severe immunodeficiency and increased infection rates (5).

Cells convert vitamin A into several related compounds, collectively known as retinoids. These include retinol and its derivative retinoic acid (RA). RA is a potent regulatory molecule that controls gene expression through RA receptors (RARa, β, and γ), members of the nuclear receptor family that activate the transcription of specific target genes (6). The intestinal epithelium plays a central role in retinoid metabolism by absorbing dietary vitamin A and expressing RA-generating enzymes and RARs (7, 8). This suggests that the epithelium could help to regulate vitamin A-dependent adaptive immunity. A recent study implicated epithelial RARa in the development of the epithelial barrier as well as lymphoid follicles that support intestinal immune cell development (8). However, it is not clear whether epithelial cells regulate the development of specific vitamin A-dependent immunological pathways including the development of gut-homing CD4^+^ T cells, IgA-producing B cells, and Th17 cell effector function.

Serum amyloid A proteins are a family of immunoregulatory proteins that highlight the integration of dietary and microbiota signals by the intestinal epithelium. SAAs are retinol-binding proteins that are expressed at the site of retinoid uptake (intestinal epithelium) and retinoid storage (liver) and circulate retinol following systemic bacterial exposure (9). In the intestine, SAAs stimulate IL-17 expression by Th17 cells (10), thus shaping their effector functions. Expression of SAAs in the intestine and the liver requires both a microbial signal (microbiota colonization in the intestine and systemic bacterial challenge in the liver) (9, 10) and dietary vitamin A (9). The microbiota triggers intestinal epithelial SAA expression through a multicellular signaling circuit involving dendritic cells, innate lymphoid cells, and epithelial STAT3 (10). However, the mechanisms by which vitamin A regulates SAA expression are unknown.

Here we show that RARβ activates *Saa* expression through direct binding to retinoic acid response elements (RAREs) in *Saa* promoters. Consistent with the known role of SAAs in regulating Th17 cell effector function (10), we also find that epithelial RARβ regulates IL-17 production by Th17 cells. More generally, we show that epithelial RARβ regulates other known vitamin A-dependent adaptive immune responses, including the development of gut-homing CD4^+^ T cells and IgA-producing B cells. Our findings thus provide insight into how the intestinal epithelium senses dietary vitamin A status to control vitamin A-dependent adaptive immunity.

## Results

### RARβ directs retinoid-dependent SAA expression

Dietary vitamin A is required for SAA expression in mouse intestine and liver, and the vitamin A derivatives retinol and RA stimulate *SAA1* and *SAA2* expression in the human liver cell line HepG2 ((9); Fig. 1*A*). To determine if retinoids also stimulate SAA expression in intestinal epithelial cells, we studied the mouse intestinal epithelial cell line MODE-K. MODE-K cells do not express *Saa1* and Saa2; however, they do express *Saa3*, which is expressed in the intestine but not the liver. Saa3 expression increased with the addition of bacterial lipopolysaccharide (LPS) and retinol (Fig. 1*B*), showing that Saa3 expression in MODE-K cells is highest in the presence of both a bacterial signal and a retinoid.

**Figure 1.**
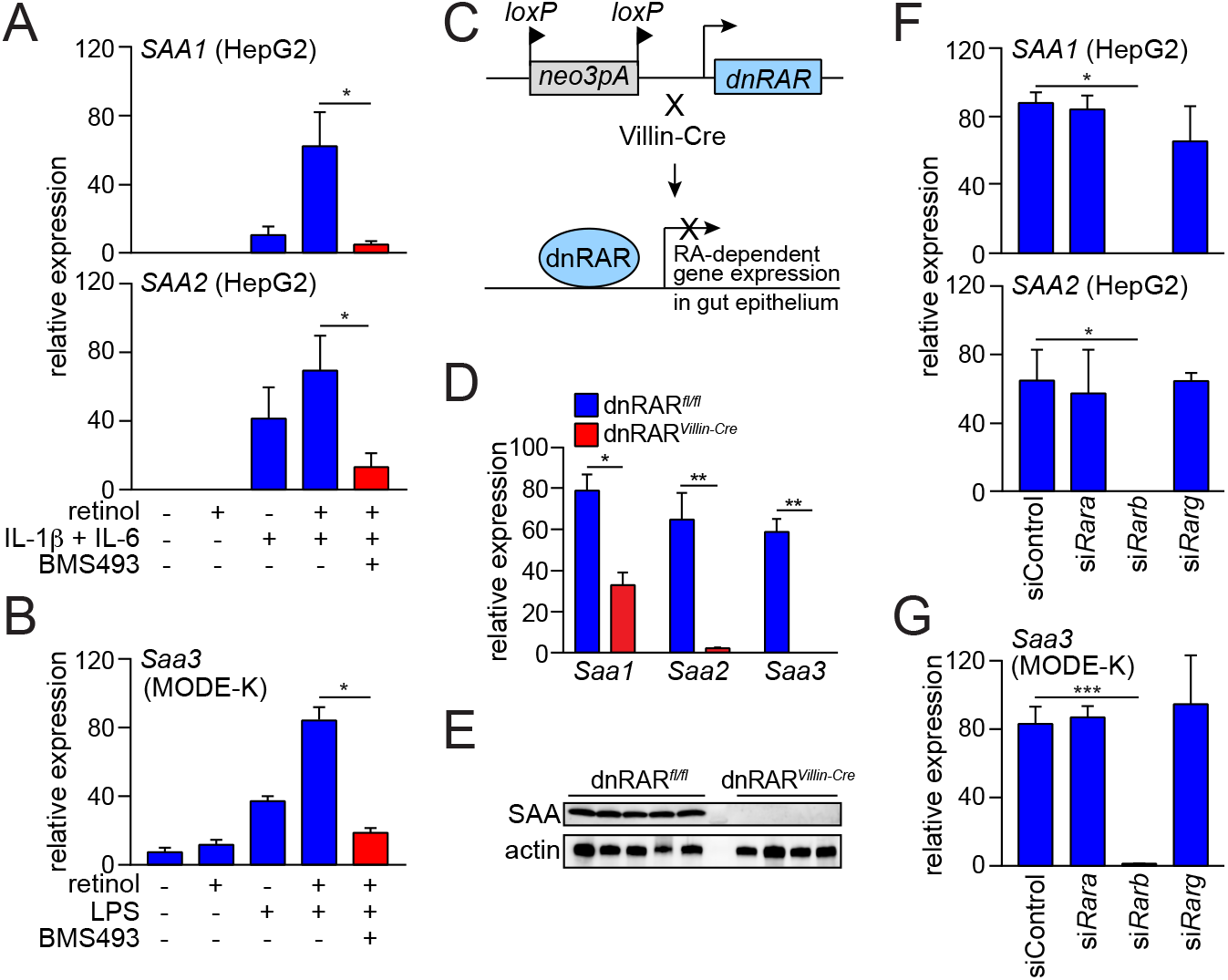
RARβ directs retinoid-dependent SAA expression. (*A,B*) qPCR analysis of *SAA*(human) and *Saa* (mouse) expression. (*A*) *SAA1 and SAA2* expression in HepG2 cells treated for 24 h with retinol, IL-1β, IL-6, and the RAR inhibitor BMS493. (*B*) *Saa3* expression in MODE-K cells treated for 24 h with retinol, LPS, and BMS493. *N*=4 replicates/group; data represent two independent experiments. (*C*) Selective disruption of RAR activity in intestinal epithelial cells. Knock-in mice carry a neomycin resistance gene and 3 loxP-flanked polyadenylation sequences that are located upstream of an open reading frame encoding a dominant negative (dn)RAR. Breeding to *Villin-Cre* mice results in selective expression of the dnRAR in intestinal epithelial cells. (*D*) qPCR analysis of *Saa* transcripts in the distal small intestines of dnRAR*^fl/fl^* mice and dnRAR^Villin-Cre^ mice. *N*=4 mice/group; data represent two independent experiments. (*E*) Western blot of SAA in the distal small intestines of dnRAR*^fl/fl^* mice and dnRAR*^Villin-Cre^* mice. (*F,G*) qPCR analysis of *SAA* (human) and *Saa* (mouse) expression after siRNA knockdown of specific *Rar* isoforms. (*F*) *SAA1* and *SAA2* were analyzed in HepG2 cells treated with retinol, IL-1β and IL-6 as in *A*, and (*G*) Saa3 expression was analyzed in MODE-K cells treated with retinol and LPS as in *B*. *N*=3 replicates/group; data represent two independent experiments. Means±SEM are plotted. \**P*<0.05, \*\**P*<0.01, \*\*\**P*<0.001 as determined by Student’s *t*-test.

To further our mechanistic understanding of how retinoids stimulate SAA expression, we tested whether RARs are required. We added the RAR inhibitor BMS493 to HepG2 cells in the presence of retinol and the cytokines IL-1β and IL-6, which are generated during systemic infection (11). We chose to use retinol instead of retinoic acid, as retinol is more stable than retinoic acid (12), freely diffuses across membranes (13), and both HepG2 and MODE-K cells convert retinol to retinoic acid (14). BMS493 inhibited retinol-dependent *SAA1* and *SAA2* expression in HepG2 cells (Fig. 1*A*) and Saa3 expression in MODE-K cells (Fig. 1*B*), suggesting that RARs are required for retinoid-induced SAA expression in cells.

To determine if RARs govern SAA expression *in vivo* we studied mice with selective disruption of RAR activity in intestinal epithelial cells. The mice were derived from knock-in mice carrying three *loxP*-flanked polyadenylation sequences upstream of an open reading frame encoding a dominant negative form of human RARa (dnRAR) that disrupts RAR activity (15, 16). We crossed mice carrying the epithelial cell-restricted *Villin-Cre* transgene (17) with the *loxP*-flanked dnRAR knock-in mice (dnRAR*^fl/fl^*) to selectively disrupt RAR activity in epithelial cells (Fig. 1*C*). Expression of *Saa1-3* and SAA protein levels were lower in the dnRAR*^Villin-Cre^* mice as compared to dnRAR controls (Figs. 1*D* and *E*), indicating that RAR activity regulates intestinal SAA expression *in vivo*.

The mouse genome encodes three RAR isoforms (RARα, β, and γ), and we therefore sought to identify which isoform governs *Saa* expression. We used small interfering (si) RNAs to target individual *Rar* isoforms in HepG2 (Fig. S1) and MODE-K cells (Fig. S2). siRNA knockdown of *Rarb* suppressed *SAA1* and *SAA2* expression in HepG2 cells (Fig. 1*F*) and *Saa3* expression in MODE-K cells (Fig. 1*G*), while knockdown of *Rara* and *Rarg* had little effect. Thus, RARβ is uniquely required for retinol-dependent *Saa* expression in cells.

### RARβ activates *Saa3* transcription by binding directly to its promoter

We next asked whether RARβ regulates SAA expression through direct binding to *Saa* promoters. RARs bind to canonical promoter sequences called retinoic acid response elements (RAREs) that consist of the direct repetition of two core motifs. Most RAREs are composed of two hexameric motifs, 5’-(A/G)G(G/T)TCA-3’, arranged as palindromes, direct repeats, or inverted repeats (18).

*In silico* analysis of the ~4.1 kb mouse Saa3 promoter region using NUBIScan (19) identified multiple potential RAREs. We selected the 30 RAREs identified by NUBIScan as having the highest statistical chance of being functional RAREs and performed chromatin immunoprecipitation (ChIP) assays for RARβ binding at each (Fig. 2*A*; Table S1). RARβ bound to the *Saa3* promoter in MODE-K cells at multiple RAREs, including those located at −224, −327, and −1740 (Fig. 2*A*). We further verified binding of RARβ to RARE −224 (Fig. 2*B*) and showed promoter activity for the 4.1 kb region by a luciferase reporter assay (Fig. 2*C*). Introduction of point mutations into RARE −224 abolished reporter expression (Fig. 2*A* and *C*), establishing that this RARE is essential for Saa3 promoter activity. Thus, RARβ activates Saa3 transcription by binding directly to its promoter. *In silico* analysis of mouse *Saa1* and *Saa2* also identified multiple putative RAREs, suggesting that RARβ also binds directly to these promoters (Fig. S3; Tables S2 and S3).

**Figure 2.**
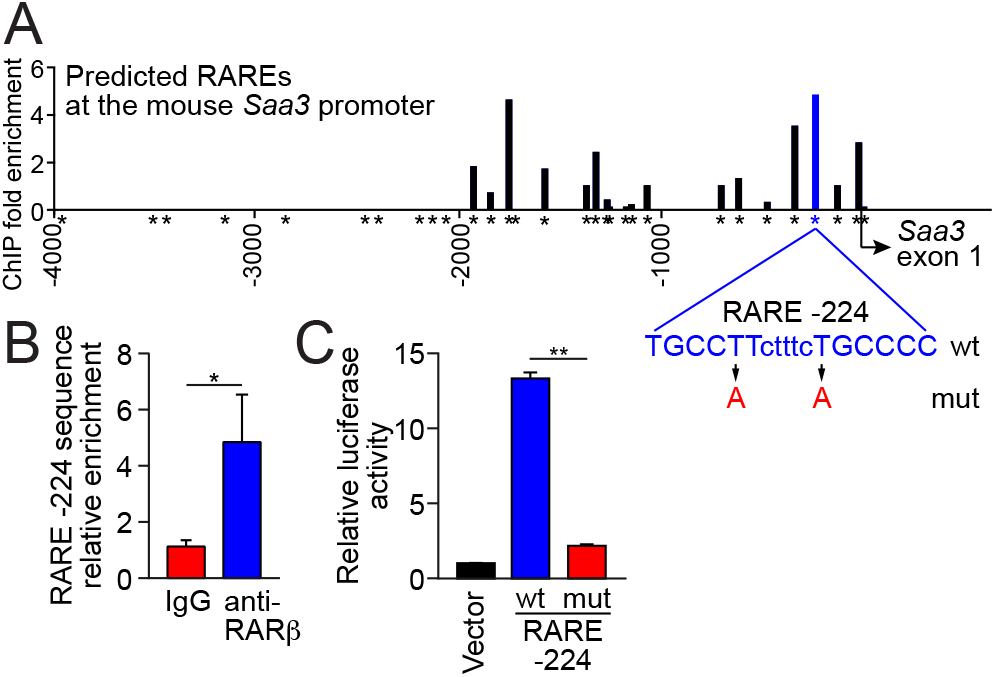
RARβ activates *Saa3* transcription by binding directly to its promoter. (*A*) RARβ binding to the Saa3 promoter was measured by chromatin immunoprecipitation (ChIP) assay with an anti-RARβ antibody. Bound promoter sequences were detected by qPCR with primers flanking each predicted retinoic acid response element (RARE, indicated by *). (*B*) The RARE located 224 nt upstream of the Saa3 start site (RARE −224) was further validated by ChIP. *N*=3 replicates per group; data represent three independent experiments. (*C*) Luciferase reporter assay for *Saa3* promoter activity. A 4103 bp fragment of the Saa3 promoter was fused to a firefly luciferase reporter and RARE −224 was mutated as shown in *A*. MODE-K cells were transfected with the wild-type (wt) or mutant (mut) reporter plasmids or empty vector and were treated with retinol and LPS for 24 h. *N*=3 replicates/group; data represent two independent experiments. Means±SEM are plotted. \**P*<0.05, \*\**P*<0.01 as determined by Student’s t-test.

### Epithelial RARβ regulates epithelial immune gene expression and promotes host resistance to intestinal bacterial infection

To determine if RARβ controls intestinal SAA expression *in vivo*, we created mice with an intestinal epithelial cell-specific deletion of *Rarb*. We crossed mice with a loxP-flanked *Rarb* allele (*Rarb^fl/fl^*) (20) with *Villin-Cre* transgenic mice (17) to produce *Rarb*^Δ*IEC*^ mice (Fig. S4). SAAs were expressed throughout the small intestinal epithelium of *Rarb^fl/fl^* mice but showed markedly reduced expression in *Rarb*^Δ*IEC*^ mice (Fig. 3*A*). qPCR analysis of laser capture microdissected epithelial cells showed reduced abundance of transcripts encoding all three mouse *Saa* isoforms (*Saa1, Saa2*, and *Saa3*) (Fig. 3*B*), and SAA protein levels were reduced in the small intestines of *Rarb*^Δ*IEC*^ mice (Fig. 3*C*).

**Figure 3.**
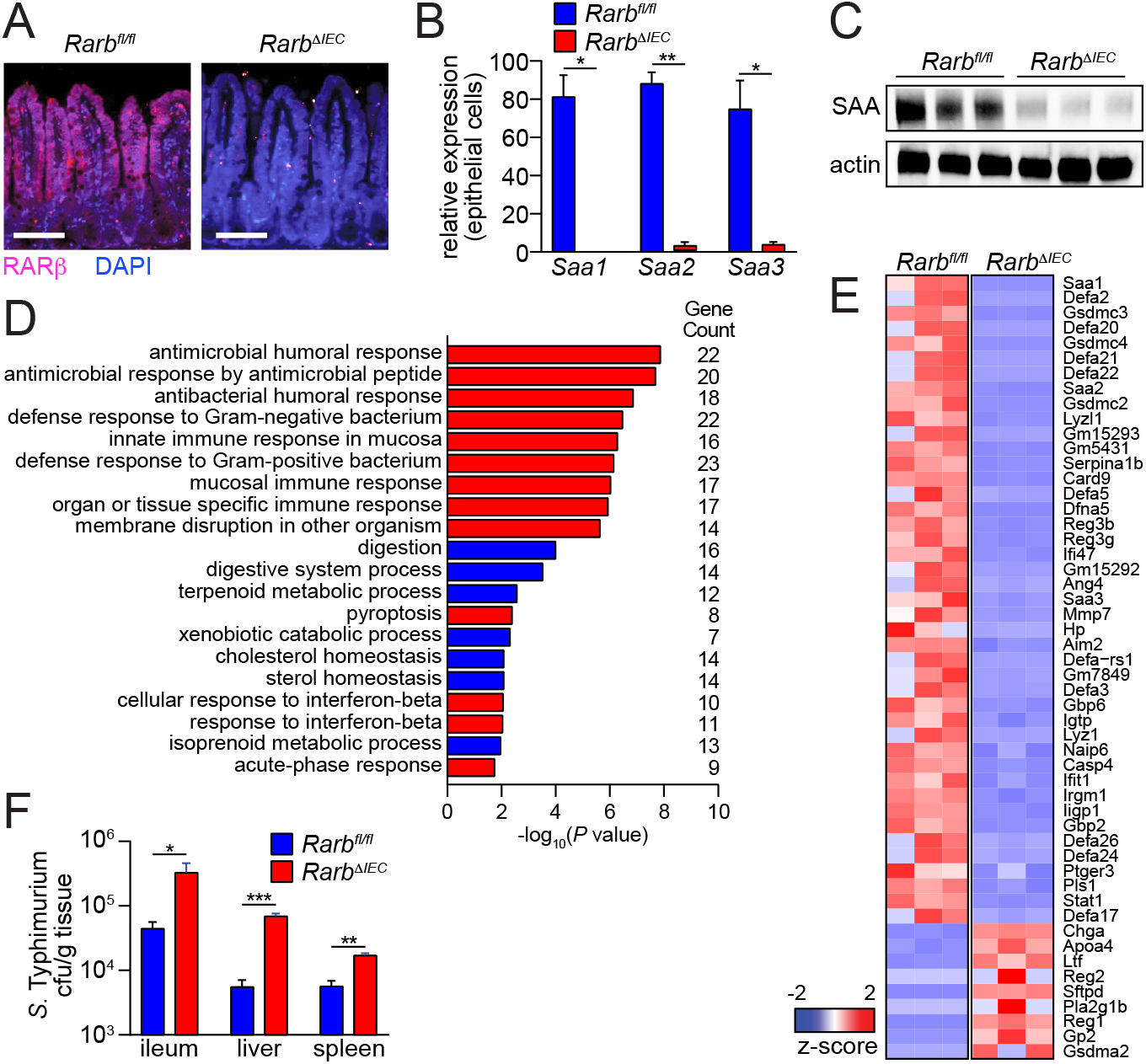
RARβ controls an epithelial immune gene expression program and promotes host resistance to bacterial infection. (*A-C*) Epithelial RARβ controls SAA expression. (*A*) Immunofluorescence detection of SAA in the small intestines of *Rarb^fl/fl^* and *Rarb*^Δ*IEC*^ mice. Scale bars, 50 μm. (*B*) qPCR analysis of *Saa* expression in small intestinal epithelial cells acquired by laser capture microdissection and (*C*) Western blot of small intestinal SAA from *Rarb^fl/fl^* and *Rarb*^Δ*IEC*^ mice, with actin as a control. (*D*) Gene ontology (GO) biological process enrichment analysis of genes identified by RNA sequencing analysis as being differentially regulated in epithelial cells from *Rarb^fl/fl^* and *Rarb*^Δ*IEC*^ mice. Immunological gene categories are highlighted in red. (*E*) Heat map displaying expression levels of the 52 genes that were identified as having immune functions by the GO analysis shown in D, and which had a −log_10_(P value) >5. (*F*) Bacterial burdens (cfu) in the distal small intestine (ileum), spleen, and liver of *Rarb^fl/fl^* and *Rarb*^Δ*IEC*^ littermates 48 hr after oral infection with 10^10^ cfu of S. Typhimurium. *N*=5 mice/group; data represent three independent experiments. Means±SEM are plotted. \**P*<0.05, \*\**P*<0.01, \*\*\**P*<0.001 as determined by Student’s t-test.

To broaden our insight into epithelial RARβ function, we identified other intestinal genes that were regulated by RARβ. We used RNAseq to compare the transcriptomes of *Rarb^fl/fl^* and *Rarb*^Δ*IEC*^ mouse small intestines, finding 832 differentially-abundant transcripts. Gene Ontology (GO) term analysis identified gene categories that were highly represented among the differentially expressed genes, including multiple categories related to immunity and metabolism (Figs. 3*D*, *E* and S5). In addition to *Saa1, Saa2*, and *Saa3* transcripts, the immunological gene category included transcripts encoding proteins involved in antimicrobial defense, including several members of the *Defa* (defensin) gene family, *Reg3b, Reg3g*, and *Ang4* (Fig. 3*E*). Also represented were transcripts encoding proteins involved in inflammasome function and assembly, including *Card9, Aim2, Naip6, Casp4*, and *Gsdmc2, 3, and 4* (Fig. 3*E*). Thus, RARβ controls an immune gene transcriptional program in intestinal epithelial cells. This suggests that epithelial sensing of vitamin A status may regulate antimicrobial defense as well as inflammasome function.

Our finding that RARβ regulates the expression of immune genes in the intestinal epithelium suggested that RARβ might promote resistance to bacterial infection of the intestine. We orally challenged *Rart*^Δ/*EC*^ and *Rarb^fl/fl^* mice with the gastrointestinal pathogen *Salmonella enterica* Serovar Typhimurium (*Salmonella* Typhimurium). 48 hours later, *Rarb*^Δ*IEC*^ mice had increased bacterial burdens in the ileum, liver, and spleen (Fig. 3*F*). The increased bacterial burdens did not arise from increased nonspecific barrier permeability (Fig. S6) or altered microbiota taxonomic composition (Fig. S7A and β), as these were similar between *Rarb*^Δ*IEC*^ and *Rarb^fl/fl^* mice. Thus, epithelial RARβ contributes to host resistance to intestinal bacterial infection and dissemination.

### Epithelial RARβ promotes intestinal Th17 cell effector function

Th17 cells are a specialized subset of CD4^+^ T cells that require the transcription factor RORγt for lineage commitment (21). Th17 cells secrete a distinct set of cytokines, including IL-17A, IL-17F, and IL-22, and promote host defense against extracellular bacteria (22). SAA secretion by intestinal epithelial cells promotes Th17 cell effector functions (10). Although SAAs are not required for Th17 cell lineage commitment, they boost Th17 cell effector function by stimulating IL-17A production in differentiated Th17 cells (10).

Our finding that epithelial RARβ governs SAA expression suggested that epithelial RARβ might also regulate IL-17 production by intestinal Th17 cells. In support of this idea, *ll17a* expression was lowered in the intestines of *Rarb*^Δ*IEC*^ mice, paralleling the lowered *ll17a* expression in *Saa1/2^−/-^* mouse intestines ((10); Fig. 4*A*). Overall frequencies of CD4^+^ RORγt^+^ Th17 cells were similar between *Rarb^fl/fl^* and *Rarb*^Δ*IEC*^ mice (Fig. 4*B*), reflecting the fact that SAA is dispensable for Th17 cell lineage commitment (10). However, IL-17A production was reduced in RORγ^+^ Th17 cells from *Rart^Δ/*EC*^ mice (*Figs. 4C* and *D*), indicating that epithelial RARβ regulates Th17 cell effector function. IL-17A production was restored by the addition of recombinant SAA1 to cultured intestinal lamina propria cells from *Rarb*^Δ*IEC*^* mice (Fig. 4*E*). This indicates that SAA1 is sufficient to rescue the defective Th17 effector function conferred by epithelial RARβ deficiency and suggests that the defect in Th17 IL-17 production in *Rart^Δ/*EC*^* mice is due to the lowered SAA expression. Thus, epithelial RARβ promotes intestinal Th17 cell effector function, likely by activating *Saa* expression.

**Figure 4.**
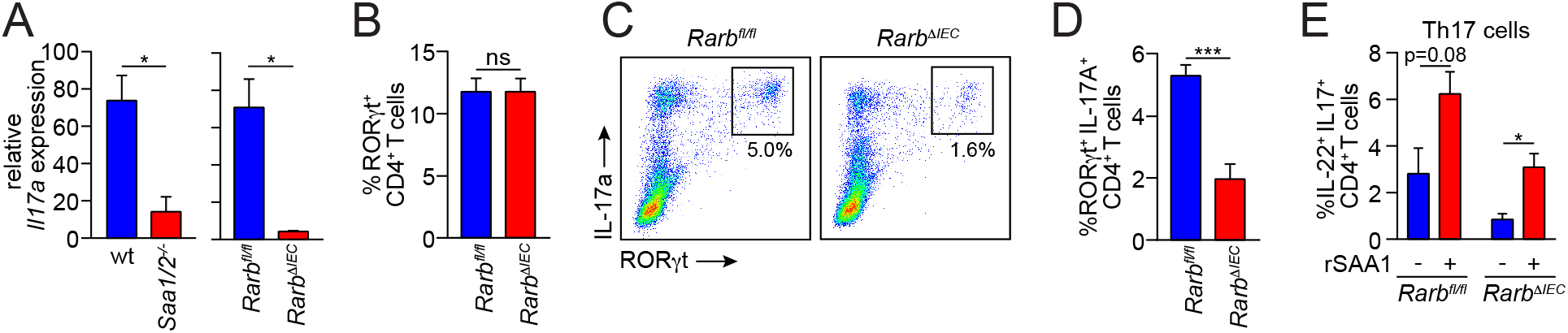
Epithelial RARβ regulates intestinal Th17 cell effector function. (*A*) qPCR analysis of *ll17a* transcripts in small intestines from wild-type (WT) and *Saa1/2^−/-^* mice, and *Rarb^fl/fl^* and *Rarb*^Δ*IEC*^ mice. (β) RORγt^+^ CD4^+^ Th17 cells as a percentage of CD45^+^ CD3^+^ cells in the small intestinal lamina propria. *N*=3 mice/group; data represent four independent experiments. (*C,D*)IL-17^+^ RORγt^+^ Th17 cells as a percentage of CD45^+^ CD4^+^ CD3^+^ cells in the small intestine. Representative flow cytometry plots (*C*) and data from multiple mice (*D*) are shown. *N*=3 mice/group; data represent four independent experiments. (*E*) Recombinant SAA1 (rSAA1) rescues lowered IL-17^+^ IL-22^+^ CD4^^+^^ T cell numbers from *Rart*^Δ/*EC*^ mice. Small intestinal lamina propria cells were isolated and stimulated *ex vivo* with ionomycin, PMA, brefeldin A, and rSAA1 for 4 hours prior to flow cytometry analysis. *N*=3 mice/group; data represent four independent experiments. Means±SEM are plotted. \**P*<0.05, \*\**P*<0.01, as determined by Student’s *t*-test. ns, not significant.

### Epithelial RARβ promotes development of gut homing T cells and IgA-producing B cells

Vitamin A and its derivative RA are essential for the development of key intestinal adaptive immune cells, including gut homing CD4^+^ T cells (3) and IgA-producing B cells (2). RA also promotes the generation of Foxp3^+^ regulatory T (T_reg_) cells (23,24). We therefore sought to determine if epithelial RARβ regulates the development of each of these cell populations. In the case of gut homing T cells, RA-producing dendritic cells (DC) migrate to the mesenteric lymph nodes (MLN), where they imprint gut homing receptors on activated CD4^+^ T cells (3). *Rarb*^Δ*IEC*^ mice had reduced frequencies of CD4^+^ T cells imprinted with the gut homing receptors CCR9 and α4β7 in the MLN and the small intestine (Figs. 5*A-C*), reduced total numbers of small intestinal CCR9^+^ α4β7^+^ CD4^+^ T cells (Fig. 5*D*), and reduced overall numbers of small intestinal CD4^+^ T cells (Figs. 5*E-G*). Thus, epithelial RARβ is essential for the development of gut homing CD4^+^ T cells.

**Figure 5.**
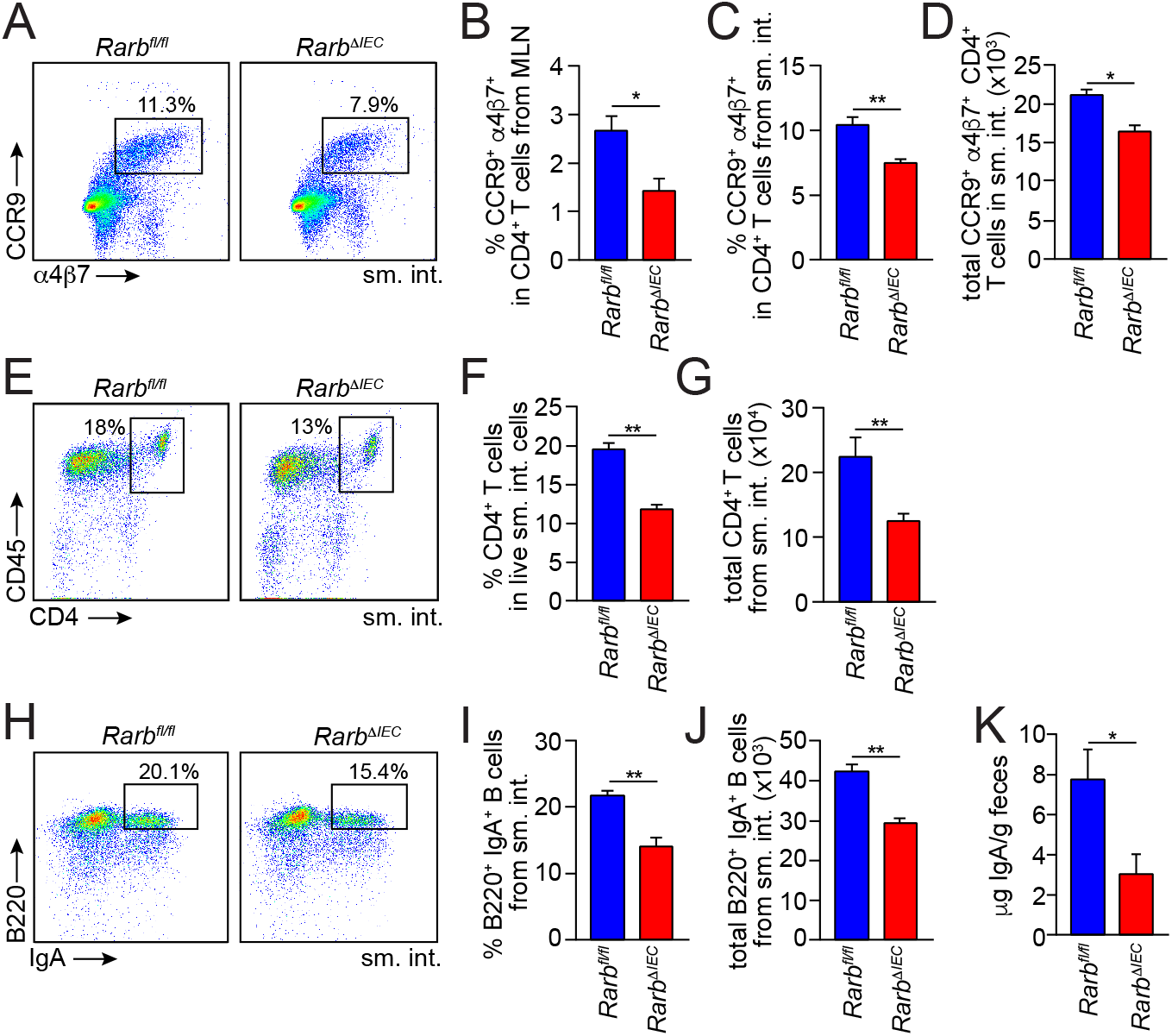
Epithelial RARβ regulates the development of gut homing T cells and IgA-producing B cells. (*A-D*) Expression of the gut homing markers a4β7 and CCR9 on T cells (CD4^+^ CD45^+^ CD3^+^) from *Rarb^fl/fl^* and *Rarb*^Δ*IEC*^ littermates. (*A*) Representative flow cytometry plots of small intestinal T cells. Frequencies of CCR9^+^ α4β7^+^ cells in CD4^+^ CD45^+^ CD3^+^ cells from MLN (*B*) and small intestine (*C*). Total gut homing CD4^+^ T cell numbers (CCR9^+^ α4β7^+^ CD4^+^ CD45^+^ CD3^+^) are given in (*D*). *N*=3 mice/group; data represent four independent experiments. (*E*) Flow cytometry of CD4^+^ (CD45^+^ CD3^+^) T cells from the small intestines of *Rarb^fl/fl^* and *Rarb*^Δ*IEC*^ littermates. CD4^+^ (CD45^+^ CD3^+^) T cell frequencies are quantified in (*F*) and total small intestinal CD4^+^ T cell numbers (CD4^+^ CD45^+^ CD3^+^) are given in (*G*). *N*=3 mice per group; data represent four independent experiments. (*H*) Flow cytometry analysis of IgA^+^ B220^+^ (CD45^+^ CD19^+^) B cells from *Rarb^fl/fl^* and *Rarb*^Δ*IEC*^ littermates. IgA^+^ B220^+^ cell frequencies in CD45^+^ CD19^+^ B cells are quantified in (*I*) and total numbers of small intestinal IgA^+^ B cells are shown in (*J*). *N*=3 mice per group; data represent four independent experiments. (*K*) Quantification of fecal IgA by ELISA. *N*=4 mice/group; data represent two independent experiments. Means±SEM are plotted. \**P*<0.05, \*\**P*<0.01 as determined by Student’s t-test. sm. int., small intestine. MLN, mesenteric lymph nodes.

RA-producing DCs also induce IgA expression in gut homing B lymphocytes (2). *Rarb*^Δ*IEC*^ mice had reduced frequencies and numbers of small intestinal IgA+ B cells (Figs. 5*H-J*) and decreased fecal IgA concentrations (Fig. 5*K*), indicating that epithelial RARβ promotes the development of IgA-producing B cells. Although DC-produced RA also promotes the development of intestinal Foxp3+ Treg cells (23,24), frequencies of Foxp3^+^ cells among small intestinal CD4^+^ T cells were similar between *Rarb^fl/fl^* and *Rarb*^Δ*IEC*^ mice (Fig. S8A and B), indicating that epithelial RARβ is not required for the generation of intestinal T_reg_ cells.

RARα is a closely related RAR isoform that impacts several aspects of intestinal immunity, including Paneth and goblet cell development, numbers of RA-producing DCs, and overall B cell numbers (8). We considered whether RARα and RARβ might have overlapping functions in the control of intestinal adaptive immunity. We found that *RARb^fl/fl^* and *Rarb*^Δ*IEC*^ mice had similar numbers of small intestinal Paneth cells or goblet cells (Fig. S9A), and similar frequencies of CD11c^+^ CD103^+^ cells (which include RA-producing DCs) (Fig. S9β) and B220+ B cells (Fig. S9C). In contrast to *Rarb*^Δ*IEC*^ mice, *Rara*^Δ/*EC*^ mice had elevated intestinal *Saa* expression (Fig. S10A and β). Numbers of intestinal gut homing T cells (Fig. S11A-D), and total CD4^+^ T cells (Fig. S11*E-G*) were also elevated relative to *Rara^fl/fl^* mice. Frequencies of intestinal IgA-producing B cells (Fig. S11H-J), and fecal IgA quantities (Fig. S11K) were similar as compared to *Rara^fl/fl^* mice. These data indicate that RARa is dispensable for CD4^+^ T cell homing and B cell expression of IgA, and show that epithelial RARa and RARβ regulate distinct aspects of intestinal immunity.

## Discussion

The intestinal epithelium is a vital interface between the environment and the underlying immune system. Epithelial cells sense key environmental factors, including the microbiota and the host diet, and use these cues to orchestrate adaptive immunity in subepithelial tissues. The *Saa* genes highlight the environmental sensing function of the gut epithelium by requiring both microbiota and vitamin A for expression in the intestinal epithelium. We have now unraveled the molecular basis for how vitamin A directs SAA expression by showing that RARβ activates the expression of *Saa* genes through direct promoter binding. More generally, we show that epithelial RARβ regulates the function of Th17 cells, the development of gut homing T cells, and the development of IgA-producing B cells. Our findings thus provide important mechanistic insight into how the intestinal epithelium senses vitamin A to regulate intestinal adaptive immunity.

The three RAR isoforms (α, β, and γ) are conserved across species (25), suggesting a unique conserved function for each isoform. Supporting this idea, we found that RARβ was uniquely required for expression of *Saa* genes, and that RARa and RARβ have non-redundant functions in the regulation of intestinal immune function. While RARa regulates the development of intestinal epithelial cell secretory lineages, RA-producing DCs, and overall B cell populations (8), our findings show that RARβ regulates Th17 cell effector function, the development of gut homing T cells, and IgA-producing B cells. Thus, epithelial RARa and RARβ are both essential for intestinal immune homeostasis but regulate distinct aspects of immunity.

Our finding that epithelial RARβ promotes IL-17 production by Th17 cells suggests that dietary vitamin A promotes Th17 cell effector function. This idea is supported by prior work showing that mice fed a vitamin A-deficient diet exhibit lowered intestinal IL-17 production (4). Furthermore, mice carrying a T cell-specific *Rara* deletion show lowered IL-17 production by Th17 cells (4), indicating that T cell-intrinsic RA signaling is required for Th17 cell effector function. Taken together, these findings indicate that Th17 cell effector function is a component of vitamin A-dependent intestinal immunity.

We propose that the function of SAAs in retinol transport could explain the requirement for both epithelial cell-intrinsic RARβ and T cell-intrinsic RARa in Th17 cell effector function. Given that production of IL-17 by Th17 cells requires vitamin A (4) and that SAAs transport the vitamin A derivative retinol, it is possible that SAAs deliver retinol directly to Th17 cells for conversion to RA and activation of RARα. Alternatively, SAAs could deliver retinol to antigen presenting cells (such as dendritic cells or macrophages) for conversion to RA and delivery to Th17 cells. Defective Th17 cell function could also help explain the increased susceptibility of *Rarb*^Δ*IEC*^ mice to S. Typhimurium infection, which require Th17 cell responses for effective clearance.

SAA could also in part account for the essential role of epithelial RARβ in the development of gut-homing T cells and IgA-producing B cells. Development of both groups of cells requires RA-producing DCs. These DCs convert retinol to RA, which imprints gut homing receptors on T cells and induces IgA expression in B cells (2, 3). Given the retinol transport function of SAAs, we propose that SAAs could deliver retinol from the epithelium to RA-producing DCs to serve as substrate for RA production. Future work will be directed at testing this idea.

Altogether, our findings provide new insight into how the intestinal epithelium uses dietary cues to orchestrate adaptive immunity in the intestine (Fig. S12). By showing that epithelial RARβ is a key regulator of vitamin A-dependent immunity, we highlight epithelial RARs as potential therapeutic targets for the modulation of intestinal immunity during infection or inflammation.

## Methods

Full methods are presented in ***SI Methods***

### Mice

*Rarb*^Δ*IEC*^ mice were generated by crossing *Rarb^fl/fl^* mice (20) with *Villin-Cre* mice (Jackson Laboratories), which express *Cre* recombinase under the control of the intestinal epithelial cell-specific *Villin* promoter (17). *dnRAR^Villin-Cre^* mice were generated by crossing dnRAR^fl/fl^ mice (16) with *Villin-Cre* mice. *Saa1/2^−/-^* mice were obtained from F. de Beer (26); RARa*^fl/fl^* mice were from P. Chambon (27) through Y. Belkaid (National Institutes of Health). 6-14 week-old mice were used for all experiments. Because microbiota composition is known to impact intestinal immune cell frequencies, we used age- and sex-matched littermates that were co-caged to minimize microbiota differences in all experiments. Experiments were performed in accordance with protocols approved by the Institutional Animal Care and Use Committees of the University of Texas Southwestern Medical Center.

### Cell Culture

The MODE-K cell line was provided by Dominique Kaiserlian (28). HepG2 cells were purchased from the ATCC. Cells were cultured in 1X DMEM with GlutaMAX, 10% FBS, 1X Penstrep, and 1X sodium pyruvate. Cells were maintained in 5% CO2 at 37°C. Prior to addition of retinol and LPS or cytokines, or siRNA treatment, cells were cultured in a serum-free medium for 48 hours. MODE-K cells were treated for 24 hours with retinol (100 nM; Sigma) and lipopolysaccharide (100 ng/ml; Sigma). HepG2 cells were treated for 24 hours with retinol (100 nM), IL-1β (50 pg/ml; R&D Systems), and IL-6 (100 pg/ml; R&D Systems).

### RNA sequencing and data analysis

RNA was extracted and purified from ileums of 3 mice per experimental group. Libraries were prepared from the RNAs and sequenced on an Illumina HiSeq 2500. Altered expression was defined as a >2-fold increase or decrease in average FPKM reads compared between the two groups. Further details are given in *SI Materials and Methods*.

### Western Blots

Total protein was isolated from cells or homogenized tissues as previously described (29). Blots were probed with antibodies against SAA (9), RARa (Affinity Bioreagents), RARβ (Invitrogen), RARg (ThermoFisher), and actin (ThermoFisher). The anti-SAA antibody was generated by Cocalico and recognizes all SAA isoforms (9).

### IgA ELISA

Total protein was isolated by homogenizing mouse feces in PBS with protease inhibitor cocktail (Roche). Samples were rotated at 4°C for 4 hours and centrifuged at 16,000g for 15 minutes. Supernatants were serially diluted to quantify IgA levels per manufacturer instructions (Invitrogen).

### Chromatin immunoprecipitation (ChIP) assays

MODE-K cells were crosslinked in PBS with 1% formaldehyde for 3 minutes at room temperature and quenched in 125 mM glycine at 4°C for 10 min. Nuclei from fixed cells were pelleted and used for chromatin immunoprecipitation per manufacturer’s instructions (Diagenode). Each immunoprecipitation reaction included chromatin from 5 × 10^6^ cells, 5 μg of goat anti-RARβ (Santa Cruz) or total goat IgG (Millipore), and 20 μl of Magna protein A beads (Millipore). Bound Saa3 promoter sequences were quantified using SYBR Green-based real-time PCR (target sequence and primers listed in Table S4). Relative enrichment of the *Saa3* promoter was calculated as the ratio of specific antibody pull-down to input DNA.

### Transcriptional reporter assays

A 4103bp (−4000 to +103) fragment of the Saa3 promoter was cloned into the pEZX-G04 vector containing two reporter genes (Genecopoieia) to generate the wild-type reporter construct. The putative RAR binding site (DR5) in the wild-type reporter construct was mutated from TGCCJTctttcTGCCCC to TGCCATctttcAGCCCC to generate the mutant reporter construct (Invitrogen GeneArt Site-Directed Mutagenesis Plus). MODE-K cells were transfected in 12-well plates (5 × 10^4^ cells/well) with 400 ng of wild-type or mutant reporter plasmid or 400 ng of empty vector. Cells were treated with retinol (100 nM) and LPS (100 ng/ml) for 24 hours. Luciferase activity was quantified using the Secrete-Pair Luminescence Assay Kit (Genecopoieia) and measured with a SpectraMax M5e plate reader (Molecular Devices). Gaussia luciferase (GLuc) activity was normalized against secreted alkaline phosphatase (SEAP) activity and then compared to the activity of cells transfected with pEZX-G04 alone.

### Lamina propria lymphocyte isolation and analysis

Small intestinal lamina propria lymphocytes were isolated as previously described (30). ~2 × 10^6^ cells were treated with 50 ng/ml phorbol myristate acetate (PMA), 1 mM ionomycin, and 1 mg/ml brefeldin A for 4 hours. For the rescue of IL-17 production, 5 μg/mL recombinant mSAA1 (R&D Systems) was added to whole lamina propria samples during the PMA/ionomycin/brefeldin A stimulation. Cells were fixed and permeabilized for 30 minutes and stained with commercial antibodies from Biolegend, BD, and eBiosciences. Flow cytometry was performed using the LSRII and data were analyzed with FlowJo software (TreeStar).

### *Salmonella* infection

*Salmonella enterica* serovar Typhimurium (SL1344) was grown in Luria-Bertani broth with ampicillin (100 μg/ml) at 37°C. Mice were infected intragastrically by gavage with S. Typhimurium at 10^10^ colony-forming units (cfu) per mouse. cfu in the small intestine, liver, and spleen were determined by dilution plating on Luria broth plates containing ampicillin (100 μg/ml).

### Statistical Analysis

All statistical analyses were performed using two-tailed Student’s t-test. *P ≤ 0.05; \*\**P* ≤ 0.01; \*\*\**P* ≤ 0.001; and ns, *P* > 0.05.

## Acknowledgements

We thank Dr. P. Chambon for the kind gift of the *Rara^fl/fl^* and *Rarb^fl/fl^* mice. This work was supported by NIH Grant R01 DK070855 (to L.V.H.), Welch Foundation Grant I-1762 (to L.V.H.), the Walter M. and Helen D. Bader Center for Research on Arthritis and Autoimmune Diseases (L.V.H.), and the Howard Hughes Medical Institute (L.V.H.). S.G. was supported by NIH T32 AI007520, and T.A.H. was supported by the Burroughs Wellcome Foundation Minority Enrichment Program and the Dermatology Foundation. RNAseq data have been deposited in the Gene Expression Omnibus repository with accession number GSE122471.

