## Supplementary Information for "Epithelial retinoic acid receptor β regulates serum amyloid A expression and vitamin A-dependent intestinal immunity"

**This PDF file includes:**

Extended Materials and Methods
Figs. S1 to S12
Tables S1 to S6
References for SI reference citations

### Extended Materials and Methods

**Quantitative PCR.** Total RNA was isolated from cells or homogenized tissues using the RNeasy Lipid Tissue Kit (Qiagen). cDNAs were synthesized using MMLV reverse transcriptase (ThermoFisher). Expression of mouse *Il17a* and human *RAR* genes were analyzed using Taqman gene expression qPCR assays (Invitrogen). Mouse *Saa* and human *SAA* genes were analyzed using SYBR Green qPCR reagents and specific primers (Table S5). Relative expression values were determined using the comparative $C_t$ ($\Delta\Delta C_t$) method, and transcript abundances were normalized to *Gapdh* transcript abundance.

**siRNA knockdown.** MODE-K or HepG2 cells cultured in a serum-free medium for 48 hours were trypsinized and plated at $5 \times 10^4$ cells per well of a 12-well cell culture plate. OptiMEM transfection reagent (Invitrogen) was combined with diluted siRNA pools (sequences are listed in Table S6) then added to freshly split cells. After gentle mixing, additional experimental treatments were added. MODE-K cells were treated for 24 hours with retinol (100 nM; Sigma) and lipopolysaccharide (100 ng/ml; Sigma). HepG2 cells were treated for 24 hours with retinol (100 nM), IL-1$\beta$ (50 pg/ml; R&D Systems), and IL-6 (100 pg/ml; R&D Systems). Cells were harvested 24 hours post-treatment.

**Laser capture microdissection and RNA purification.** Laser capture microdissection was performed as described (1). In brief, 5 cm of distal small intestine was washed and snap-frozen in optimum cutting temperature (OCT) compound (Fisher 23-730.571). Frozen sections were cut to a thickness of 7 µm and fixed in 70% ethanol, then stained with Methyl Green and eosin. Freshly stained sections were immediately used for laser capture microdissection of intestinal epithelial cells using an Arcturus PixCell IIe system, and 5,000-10,000 pulses were obtained from each section. RNA was extracted from isolated intestinal epithelial cells by incubating with 14 μl RNA Extraction Buffer from the PicoPure RNA Isolation Kit (Life Technology 12204-01) for 30 min, then stored at -80°C until use. Extracted RNA was purified using PicoPure RNA Isolation Kit following the manufacturer's protocol. RNA quality and concentration were determined by Agilent 2100 Bioanalyzer or RiboGreen S2 RNA Assay Kit (Thermo Fisher R11490).

**RNA sequencing (RNAseq) and data analysis.** RNA was extracted and purified from ileums of 3 mice per experimental group. RNA quality was assessed on an Agilent 2100 Bioanalyzer. Sequencing libraries were prepared using the TruSeq RNA sample preparation kit v2 (Illumina) and sequenced on an Illumina HiSeq 2500 for signal end 50 bp length reads. Sequence data were mapped against the mm10 genome using TopHat (2) and FPKMs were generated using Cuffdiff (3) with default parameters. Altered expression was defined as a >2 fold increase or decrease in average FPKM reads compared between the two groups.

**RNAseq Ontology Analysis.** RNAseq identified 832 genes were differentially expressed, greater than 2-fold, and with *p* values <0.05. These 832 genes were analyzed with the PANTHER Gene Ontology Classification System to identify enriched GO Biological Processes. The top 20 differentially regulated GO Biological Processes (gene categories) are shown in Fig. 3D, organized as immunity-related and metabolism-related genes. Bonferroni correction for multiple testing and a *p* value filter of <0.05 were applied to the PANTHER analysis.

**Retinoid Quantification.** Retinol and retinoic acid (Sigma) were freshly reconstituted in ethanol and quantified under red or amber lighting. Quantification based on absorbance at 325 nm (retinol) and 350 nm (retinoic acid) is in accordance with previously published molar extinction coefficients (4). Retinoids for use in cell culture treatments were utilized in a vehicle of 0.1% ethanol in cell culture medium.

**Flow Cytometry Reagents and Antibodies.** Foxp3/transcription factor staining reagents for fixation and permeabilization were used (eBiosciences). Zombie Yellow (Biolegend) fixable viability dye was used to gate live cells prior to further analysis. Antibodies used to characterize Th17 populations include: CD45-FITC (Biolegend), CD4-PECy7 (Biolegend), CD3-APC-Cy7 (Biolegend), RORgt-PE (eBiosciences), IL-17a-BV421 (BD Bioscience), IL-22-APC (eBiosciences). Antibodies used to characterize gut homing T cells include: CD3-APC (R&D Systems), CD45-FITC (Biolegend), CD4-PECy7 (Biolegend), CCR9-PE (Biolegend), $\alpha 4\beta 7$-Pacific Blue (BD Bioscience). Antibodies used to characterize Foxp3 T regulatory (Treg) cells include Foxp3-PE-Cy5 (eBiosciences), CD45-FITC (eBiosciences), CD4-Brilliant Violet 421 (Biolegend), CD3-PE (eBiosciences). Antibodies used to characterize IgA-producing B cells include: CD45-BV650 (Biolegend), CD19-FITC (eBiosciences), B220-APC (eBiosciences), and IgA-PE (eBiosciences). Antibodies used to characterize dendritic cell populations include CD45-Pacific Blue (Biolegend), CD11c-APC-Cy7 (eBiosciences), CD103-FITC (Biolegend), MHCII-BV711 (eBiosciences), CD11b-PE (Biolegend), and F480-APC (eBiosciences).

**16*S* rRNA gene sequencing and data analysis**

Fecal samples were collected from $Rarb^{fl/fl}$ and $Rarb^{\Delta IEC}$ littermates that were co-caged. Fecal DNA was purified using the FastDNA Spin Kit (MP Biomedicals 116560-200) and a FastPrep-24 Homogenizer. Sequencing libraries were prepared using the HotStarTaq Plus Master Mix Kit

(Qiagen) with primers flanking variable regions V3-V4. Sequencing was performed on a MiSeq following the manufacturer's guidelines. Operational taxonomic units (OTUs) were defined by clustering at 3% divergence (97% similarity). Final OTUs were taxonomically classified using BLASTn against a curated database derived from RDPII and NCBI. Principle component analysis was performed using the "prcomp" function in R and differential abundance analysis was performed using DESeq2 (5).

**Histology.** Paraffin embedded sections of Bouin's fixed mouse distal ileum were cut, and heat-induced epitope retrieval was performed in a 0.05% sodium citrate buffer (pH 6.0). The slides were incubated with rabbit anti-mouse SAA antiserum (6)(1:50 dilution) at 4°C overnight and then detected with a goat anti-rabbit IgG Cy3 conjugate (Biomeda). Tissues were counterstained with DAPI and images were captured on a Zeiss AxioImager M1 microscope. H&E and PAS staining were performed by the UTSW Histology Core.

**Supplementary Information References**

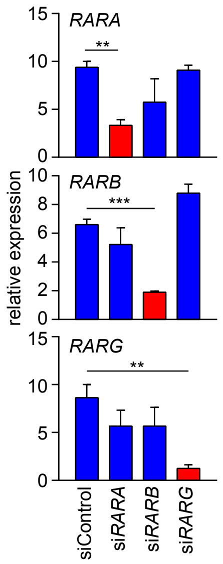


**Figure S1: siRNA knockdown of *RAR* expression in HepG2 cells.** HepG2 cells were treated for 24 hours with retinol (100 nM), IL-1β (50 pg/ml), IL-6 (100 pg/ml), and were transfected with siRNAs targeting each of the three *RAR* isoforms. A scrambled siRNA was used as a negative control (siControl). *RAR* expression was quantified by qPCR analysis. Assays were performed in triplicate and represent three independent experiments. Means±SEM are plotted. ***P*<0.01, ****P*<0.001 as determined by Student's *t*-test.

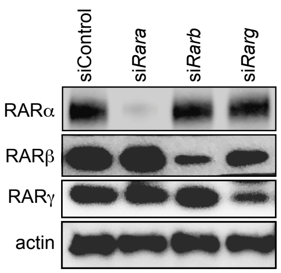


**Figure S2: siRNA knockdown of RAR expression in MODE-K cells.** MODE-K cells were treated for 24 hours with retinol (100 nM), IL-1β (50 pg/ml), IL-6 (100 pg/ml), and were transfected with siRNAs targeting each of the three *Rar* isoforms. RARα, β, and γ were detected by Western blot using antibodies specific to each isoform, with actin as a loading control. The blot is representative of three independent experiments.

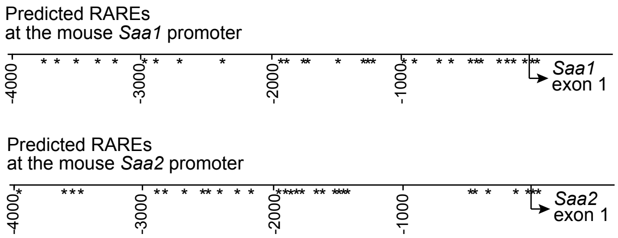


**Figure S3: Predicted retinoic acid response elements (RAREs) in the mouse *Saa1* and *Saa2* promoters.** *In silico* analysis of the *Saa1* and *Saa2* promoter regions using NUBIScan (7) identified predicted RARE sequences in each promoter.

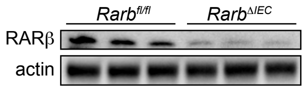


**Figure S4: Epithelial cell-specific deletion of *Rarb* in mice.** $Rarb^{\Delta IEC}$ mice were created by crossing $Rarb^{fl/fl}$ mice with mice carrying a *Villin-Cre* transgene. Epithelial cells were isolated from $Rarb^{\Delta IEC}$ and $Rarb^{fl/fl}$ mice by treatment with 10 mM EDTA as previously described (8). RARβ was detected by Western blot, with actin as a loading control.

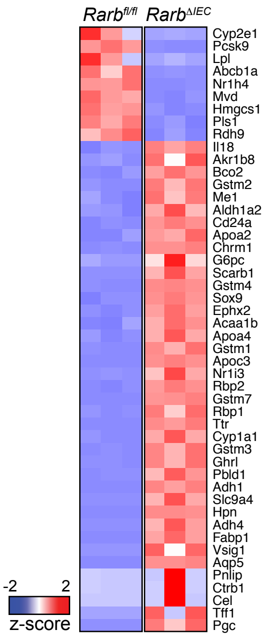


**Figure S5: Altered expression of metabolism genes in small intestines from mice with an epithelial cell-specific deletion of *Rarb*.** RNAseq analysis was performed on small intestines from *Rarb*$^{fl/fl}$ and *Rarb*$^{\Delta IEC}$ mice. Gene ontology (GO) biological process enrichment analysis of differentially regulated genes is shown in Fig. 3*D*, with metabolic gene categories highlighted in blue. Heat map shows expression levels of the 48 genes identified as having metabolic functions by the GO analysis, and which also had a $-\log_{10}(P \text{ value}) > 5$.

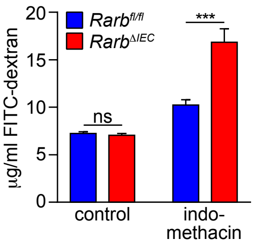


**Figure S6: Intestinal permeability measurements in *Rarb*$^{fl/fl}$ and *Rarb*$^{\Delta IEC}$ mice.** Serum concentrations of FITC-dextran 4 hours after oral gavage are shown. As a positive control, intestinal epithelial damage was induced by pretreatment with indomethacin (15 mg/kg in DMSO) for 1 hour prior to FITC-dextran administration (600 mg/kg body weight; 4 kDa; Sigma). $N$=3 mice/group, and each sample was analyzed in triplicate. Means±SEM are plotted. ***$P$<0.001 as determined by Student's $t$-test. ns, not significant.

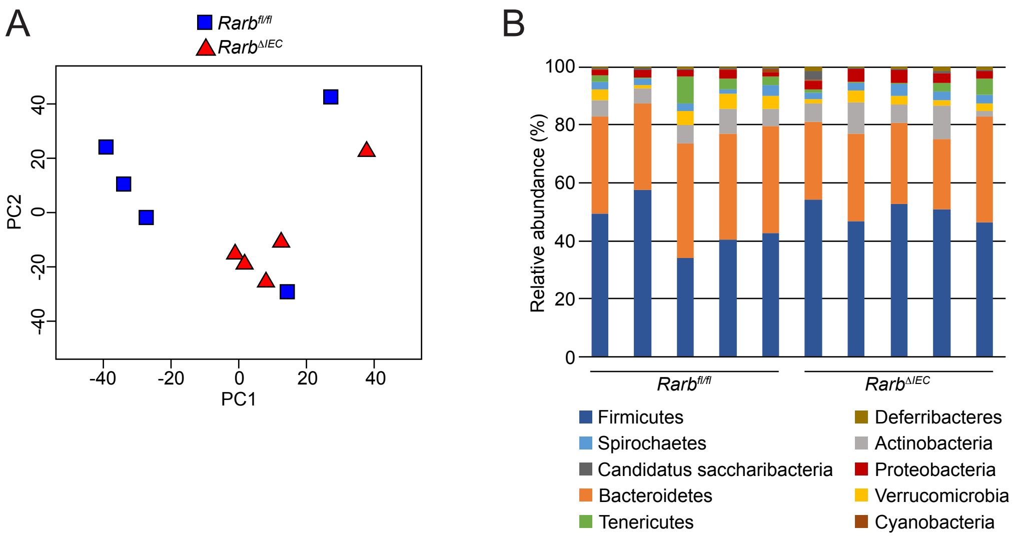


**Figure S7: The intestinal microbiotas of *Rarb*$^{fl/fl}$ and *Rarb*$^{\Delta IEC}$ mice are similar.** (*A*) Principal coordinate analysis of 16*S* rRNA sequencing of fecal samples from *Rarb*$^{fl/fl}$ and *Rarb*$^{\Delta IEC}$ mice. The mice were littermates of heterozygous crosses that remained cohoused. Each dot represents one mouse. (*B*) Relative abundances of bacterial genera in *Rarb*$^{fl/fl}$ and *Rarb*$^{\Delta IEC}$ mice.

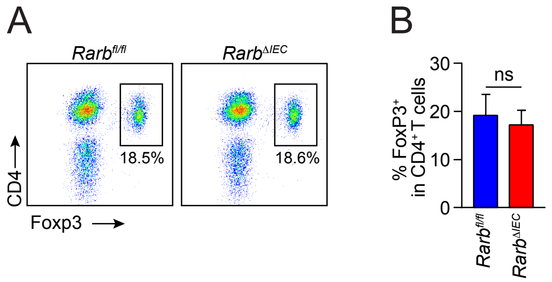


**Figure S8: $T_{reg}$ cell frequencies are similar in the intestines of *Rarb*$^{fl/fl}$ and *Rarb*$^{\Delta IEC}$ mice.** (*A*) Representative flow cytometry plots of T cells ($CD45^+$ $CD3^+$ $CD4^+$) expressing Foxp3, which marks $T_{reg}$ cells. The T cells were isolated from the small intestines of *Rarb*$^{fl/fl}$ and *Rarb*$^{\Delta IEC}$ littermates. (*B*) Quantification of $T_{reg}$ frequencies in *A*. *N*=8 (*Rarb*$^{fl/fl}$) and 6 (*Rarb*$^{\Delta IEC}$) mice per group from two independent experiments. ns, not significant as determined by Student's *t*-test.

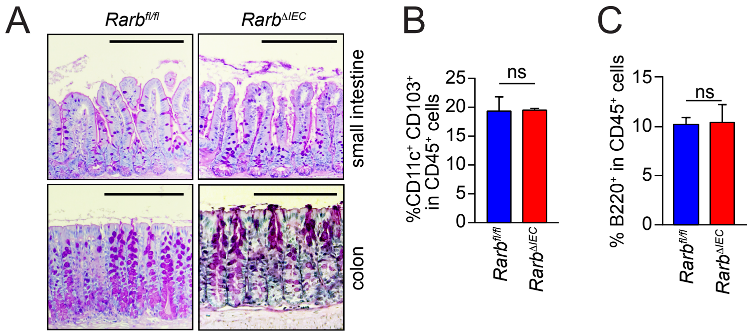


**Figure S9: Numbers of Paneth cells and goblet cells, and frequencies of $CD11c^+$ $CD103^+$ dendritic cells and B cells are similar in the intestines of *Rarb*$^{fl/fl}$ and *Rarb*$^{\Delta IEC}$ mice.** (*A*) Periodic Acid Schiff (PAS) staining was performed on small intestines and colons from *Rarb*$^{\Delta IEC}$ mice and their *Rarb*$^{fl/fl}$ littermates in order to detect goblet cells. Scale bars, 220 μm. (*B* and *C*) Cell suspensions from the small intestines of *Rarb*$^{\Delta IEC}$ mice and their *Rarb*$^{fl/fl}$ littermates were analyzed by flow cytometry. (*B*) DCs were identified using antibodies against CD45, CD11c, and CD103. Frequencies of $CD11c^+$ $CD103^+$ cells (which include RA-producing DCs) within the $CD45^+$ cell subset are shown. (*C*) B cells were identified using antibodies against CD45 and B220. Frequencies of $B220^+$ cells within the $CD45^+$ cell subset are shown. *N*=3 mice per group; data represent three independent experiments. Means±SEM are plotted. ns, not significant as determined by Student's *t*-test.

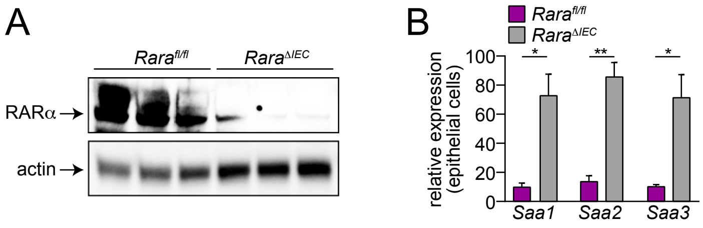


**Figure S10: Epithelial cell RARα is dispensable for *Saa* expression in the small intestine.** (*A*) *Rara*$^{\Delta IEC}$ mice were created by crossing *Rara*$^{fl/fl}$ (9) with mice carrying a *Villin-Cre* transgene (10). RARα expression was assessed by Western blotting of small intestinal epithelial cells isolated by EDTA treatment as described (8). Actin was detected as a control. (*B*) *Saa* expression was measured by qPCR analysis of small intestinal epithelial cells isolated by EDTA treatment as in *A*. *N*=3 mice per group. **P*<0.05, ***P*<0.01 as determined by Student's *t*-test.

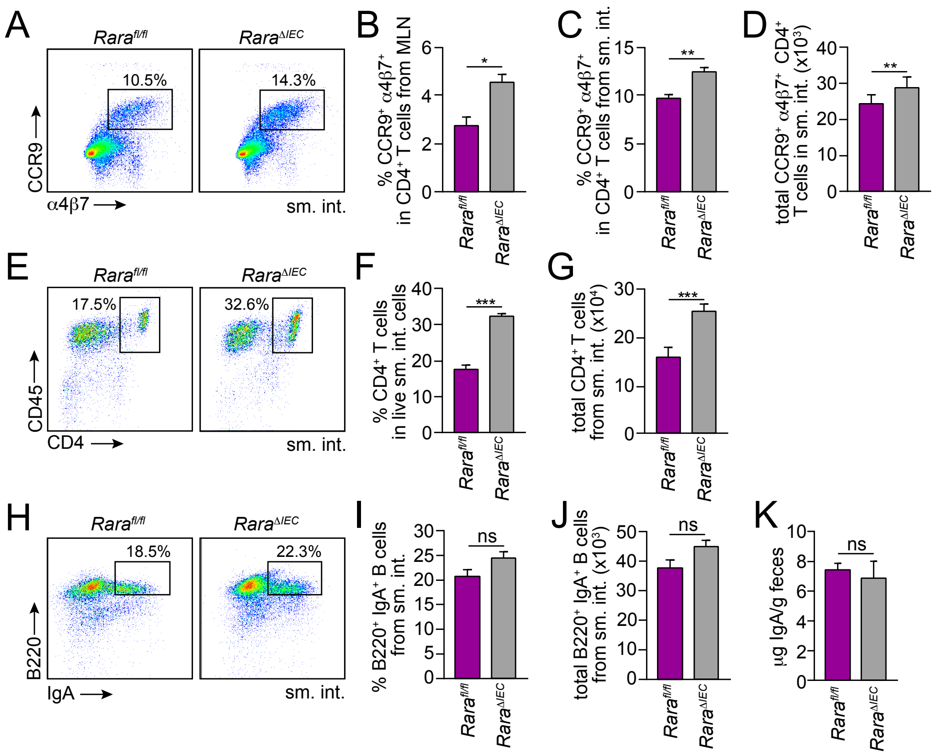


**Figure S11: Epithelial cell RARα is dispensable for $CD4^+$ T cell homing and B cell expression of IgA.** (*A*-*D*) Expression of the gut homing markers α4β7 and CCR9 on T cells ($CD4^+$ $CD45^+$ $CD3^+$) from *Rara*$^{fl/fl}$ and *Rara*$^{\Delta IEC}$ littermates. (*A*) Representative flow cytometry plots of small intestinal T cells. Frequencies of $CCR9^+$ $\alpha4\beta7^+$ cells in $CD4^+$ $CD45^+$ $CD3^+$ cells from MLN (*B*) and small intestine (*C*). Total gut homing $CD4^+$ T cell numbers ($CCR9^+$ $\alpha4\beta7^+$ $CD45^+$ $CD4^+$ $CD3^+$) are given in (*D*). N=3 mice/group; data represent three independent experiments. (*E*) Flow cytometry of $CD4^+$ ($CD45^+$ $CD3^+$) T cells from the small intestines of *Rara*$^{fl/fl}$ and *Rara*$^{\Delta IEC}$ littermates. $CD4^+$ ($CD45^+$ $CD3^+$) T cell frequencies are quantified in (*F*) and total small intestinal $CD4^+$ T cell numbers ($CD4^+$ $CD45^+$ $CD3^+$) are given in (*G*). *N*=5 mice per group; data represent two independent experiments. (*H*) Flow cytometry analysis of $IgA^+$ $B220^+$ ($CD45^+$ $CD19^+$) B cells from *Rara*$^{fl/fl}$ and *Rara*$^{\Delta IEC}$ littermates. $IgA^+$ $B220^+$ cell frequencies in $CD45^+$ $CD19^+$ B cells are quantified in (*I*) and total numbers of small intestinal $IgA^+$ B cells are shown in (*J*). *N*=5 mice per group; data represent two independent experiments. (*K*) Quantification of fecal IgA by ELISA. *N*=4 mice/group; data represent two independent experiments. Means±SEM are plotted. **P*<0.05, ***P*<0.01, ****P*<0.001 as determined by Student's *t*-test. ns, not significant. sm. int., small intestine. MLN, mesenteric lymph nodes.

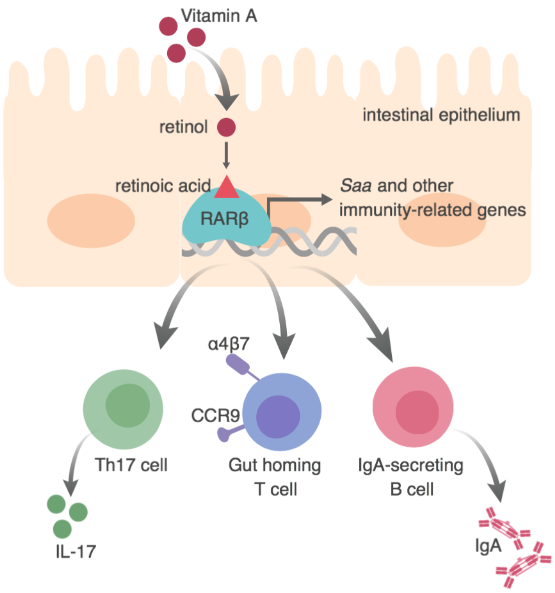


**Figure S12: Epithelial RARβ regulates intestinal adaptive immunity.** The transcription factor RARβ is activated by retinoic acid, which is derived from dietary vitamin A. Epithelial RARβ controls an immune transcriptional program that includes members of the *Saa* family. Epithelial RARβ promotes IL-17 production by Th17 cells, which is known to require SAA (11). Epithelial RARβ also promotes vitamin A-dependent immune responses that include the generation of gut homing CD4$^{+}$ T cells (marked by the gut homing receptors α4β7 and CCR9)(12) and IgA-secreting B cells (13). Our findings reveal a mechanism by which the intestinal epithelium regulates adaptive immunity in response to diet.

**Table S1: Predicted RARE sequences in the mouse *Saa3* promoter**

| Predicted RARE sequence | RARE type[1] | RARE location[2] | RARβ ChIP fold enrichment |
|---|---|---|---|
| TGAGTGgcttcTGTCCT | DR5 | -3895 | ND |
| TAAGCTaaTGCCCT | DR2 | -3556 | ND |
| AGCTCAgttcaGGTTTA | DR5 | -3537 | ND |
| TGGTCTtcTCTCCT | DR2 | -3142 | ND |
| GAGGCAatAGCTCA | DR2 | -2949 | ND |
| TTGGCTcacgtTGTCCT | DR5 | -2490 | ND |
| TGTCCTagcagTAAGCA | DR5 | -2479 | ND |
| GGTGCCtgtagAGGTCA | DR5 | -2230 | ND |
| AGGTCAgaAAATGA | DR2 | -2116 | ND |
| TTAACTgcTGACCC | DR2 | -2007 | ND |
| TGGACTttttcTTTACT | DR5 | -1915 | 1.9 |
| AGTGCAcacccAAGTCC | DR5 | -1832 | 0.8 |
| ACAACAacaacAGAGCA | DR5 | -1740 | 4.7 |
| ACAACAacAGAGCA | DR2 | -1737 | ND |
| TATCCTgttgcTGCCCT | DR5 | -1563 | 1.8 |
| AGGACAcaggaGGGACA | DR5 | -1357 | 1.1 |
| AGACAAgtgaaGGGTCA | DR5 | -1310 | 2.5 |
| TGCACAaggccTGTTCT | DR5 | -1255 | 0.5 |
| TGTTCTccTGAATG | DR2 | -1244 | 0.2 |
| CGCACAgccccTGGACA | DR5 | -1160 | 0.2 |
| TCCTCTaaggaTGCTCT | DR5 | -1134 | 0.3 |
| AGACCCagacaTGCTCT | DR5 | -1058 | 1.1 |
| TGCTCTgccccTGACTG | DR5 | -689 | 1.1 |
| TGAACAaaTTTCCT | DR2 | -603 | 1.4 |
| TGGCCAgcaagAGGTCA | DR5 | -463 | 0.4 |
| TGAGCTcacacGGGTCT | DR5 | -327 | 3.6 |
| TGCCTTctttcTGCCCC[3] | DR5 | -224 | 4.9 |
| TCACATaaggtTGCCCT | DR5 | -116 | 1.1 |
| AGGTGAtgaatAGTTAA | DR5 | -11 | 2.9 |
| TTACCTgcTGAACG | DR2 | 15 | 0.2 |

[1]RARE type: The most frequent direct repeats (DRs) with 1, 2, or 5 nucleotides spacing are termed DR1, DR2, and DR5 elements, respectively.

[2]RARE location was determined relative to the *Saa3* start site.

[3]RARE -224 was further validated by ChIP and luciferase reporter assay for Saa3 promoter activity as shown in Fig. 2*B* and *C*.

**Table S2: Predicted RARE sequences in the mouse *Saa1* promoter**

| mSAA1 | Predicted RARE sequence | RARE type | RARE location |
|---|---|---|---|
| 243 | AGGACAcaccaTGATCT | DR5 | -3758 |
| 335 | GGATCAcaacaATCCCA | DR5 | -3666 |
| 558 | AGGTCTtaGGGCCA | DR2 | -3443 |
| 693 | ACTTCAgtataGGAACA | DR5 | -3308 |
| 874 | AGAGCTagctgGGGACA | DR5 | -3127 |
| 1056 | GGATGAagAGCTCA | DR2 | -2945 |
| 1113 | AGGTCActtctTGTAAA | DR5 | -2888 |
| 1304 | AATTCAcaCGGACA | DR2 | -2697 |
| 1689 | TGTCCActaggCGGACA | DR5 | -2312 |
| 2018 | AGACCTgtagcAGGGCA | DR5 | -1983 |
| 2029 | AGGGCAgtagtAGCACA | DR5 | -1972 |
| 2234 | GGCTCAgcAGGTCA | DR2 | -1767 |
| 2330 | AGGGCAtcAGATCT | DR2 | -1671 |
| 2648 | AGACCAggccaGTCTCA | DR5 | -1353 |
| 2840 | AGAGCAagACGACA | DR2 | -1161 |
| 2863 | ATGGCAgaaggAGGAGA | DR5 | -1138 |
| 2898 | AGCAGAcaAGCTCA | DR2 | -1103 |
| 3051 | GGTTAAgaAGCACA | DR2 | -950 |
| 3182 | AGGGGAaggctGGGCCA | DR5 | -819 |
| 3304 | AGTTCCctgtgGGGCCA | DR5 | -697 |
| 3653 | AGGTAAaatggGGGGCA | DR5 | -348 |
| 3664 | GGGGCAgggggAGAACA | DR5 | -337 |
| 3664 | GGGGCAggGGGAGA | DR2 | -337 |
| 3825 | TGTGCAatgggAGCACA | DR5 | -176 |
| 3845 | GGATGAagAGCTCA | DR2 | -156 |
| 3853 | TCTTCAtccacAGGTCA | DR5 | -148 |
| 3942 | AGGACAgcCTGGCA | DR2 | -59 |
| 3953 | AGGTGAgggcaAGGACA | DR5 | -48 |
| 4003 | AGATCAccAGATCT | DR2 | 3 |
| 4031 | AGGTGAgaggcAGATCC | DR5 | 31 |

**Table S3: Predicted RARE sequences in the mouse *Saa2* promoter**

| mSAA2 | Predicted RARE sequence | RARE type | RARE location |
|---|---|---|---|
| 334 | GGTACAcaGTGTCA | DR2 | -3667 |
| 433 | AGGACTggttgAGAGCA | DR5 | -3568 |
| 522 | AGGGCAgaGGTAGA | DR2 | -3479 |
| 1130 | GGGCCAagAGAACG | DR2 | -2871 |
| 1166 | GGGTCAagagcAGCCAA | DR5 | -2835 |
| 1277 | AGCACAatagcAGGGCC | DR5 | -2724 |
| 1489 | AGATCTcaTGTGCA | DR2 | -2512 |
| 1630 | GGATCAtctcaAGTGCC | DR5 | -2371 |
| 1702 | AAGTGAgttccAGGACA | DR5 | -2299 |
| 1878 | AGATGAgagacTGGGCA | DR5 | -2123 |
| 2096 | AGTGCAgctatGGGGCA | DR5 | -1905 |
| 2148 | AAGCCActGGTCCA | DR2 | -1853 |
| 2155 | TGGTCCacaagAGGACA | DR5 | -1846 |
| 2271 | AGTACAgcAGGACT | DR2 | -1730 |
| 2428 | AAGTCAtgcagTGGTCA | DR5 | -1573 |
| 2433 | AATTCAagAGTTTA | DR2 | -1568 |
| 2471 | AGGGGAacTGGGCA | DR2 | -1530 |
| 2564 | GGGGCAgtttaATGTCA | DR5 | -1437 |
| 2572 | AGGTCTgtGGGGCA | DR2 | -1429 |
| 2583 | AGGACCctggtAGGTCT | DR5 | -1418 |
| 3413 | AGTGTAtatttTGTTCA | DR5 | -588 |
| 3421 | AGCTCAttAGTGTA | DR2 | -580 |
| 3670 | TGTTCCtgcagAGAACA | DR5 | -331 |
| 3813 | AGCACAggAGACAA | DR2 | -188 |
| 3824 | TGTGCAatgggAGCACA | DR5 | -177 |
| 3844 | GGATAAagAGCTCA | DR2 | -157 |
| 3941 | AGGACAgcCTGGCA | DR2 | -60 |
| 3958 | TGCTCAggtgaGGGGCA | DR5 | -43 |
| 4003 | AGACCAccAGATCT | DR2 | 3 |
| 4031 | AGGTGAgaggcAGATCC | DR5 | 31 |

### Table S4: PCR primers for predicted RAREs in mouse *Saa3* promoter

| Target Sequence in mSAA3 promoter | Forward Primers (5' to 3') | Reverse Primers (5' to 3') |
|---|---|---|
| RARE -3895<br>RARE -3556 | GCAAAGAGGAAATGCTGGAAAG<br>AAAGTTCTGGCAACTCACCTC | TCAAACAGGGATTGCTCCATTA<br>TAGATATCCTCGGCCACATCTC |
| RARE -3895 | GCTGCGAGCTCCTTCTG | GGGTAAGTGGGATTATGCAAGA |
| RARE -3556<br>RARE -3537 | CGCCATCCTGGTTGGAAATAA<br>TCATGGGATCCAATCCAAAGG<br>ATGCTTTATATCCAGTTTCTTGGG | CCCATCCTCCTGTTCCCTAA<br>TGCTCCCTGACCTTCCATA<br>CAGCGTTGGGTGCTAAATG |
| RARE -3142<br>RARE -2949 | CATTTGGCCTCCTGCTCTAA | CCTTCCAGCTTACGAGGTTTAT |
| RARE -3142 | TTGCCGTGGGAGTTTATCTTAT<br>CAGCTGCTAGAGCCAGAAA<br>GCAAGAGGAGACACAGGAATG | GCAGCTATACCGTCCTTCTTT<br>GTGATACCTTCACCACACCTAC<br>CCTTCTTTCTCTAGGGAACTCAATAC |
| RARE -2949 | GGTGTGGTGAAGGTATCACAT<br>GAAGCAGAGGAAAGCAGAGG<br>CCTAGAGAAAGAAGGACGGTATAG | CGAGATAGCCACAGTTCTCTTC<br>CGGGTATTATTCACCACCTTCC<br>CTTGGTAGCAAACAACTTGGG |
| RARE -2490<br>RARE -2479 | GCTGCCTTTGGCTTAGGA<br>TGGTCCTCAGCCAAGACA | AATCCCGGAACCAAATGTTTATG<br>GCCTAGGAAGTCAGAACAAAGG |
| RARE -2490 | GCTCTGTGACTGAAAGGGAAA | GGGAAGTTGTAGCTGCTTACTG |
| RARE -2479 | AGTAAGCAGCTACAACTTCCC | GTGGCTGAGGAAGTCAATACA |
| RARE -2230<br>RARE -2116 | CCTCCCTCTCGTCCCTTT<br>CCAAATGGATGCATGTGACC | CCAGAGGACCTGGGTTTAATTC<br>GTACTCATGGTAGCTCACAACT |
| RARE -2007 | CCAAATGGATGCATGTGACC | AAAGGGTAAAGGGTGGATATGAG |
| RARE -1915<br>RARE -1832 | TCTTTGATAGAAGGCAAGTGAGT | AAAGGGTAAAGGGTGGATATGAG |
| RARE -1740<br>RARE -1737 | GCAACCTTGCCCGAAATAAA<br>TCATATCCACCCTTTACCCTTTG | GTGAAGATGTGTGGACAGAAGA<br>CACAGTGACACACTCTAGCTTC |
| RARE -1563 | GTTCCTCGGAACAAGTCCATT<br>TCTTCTGTCCACACATCTTCAC<br>GTCACTGTGTCTCTCCAATGTAT<br>CAGTACCAATACTGGGTCCTTT | GGCCCAGCTCACAATCTATATC<br>TGGGATTCAAGGGACAGTTATG<br>GCTGACCCAAAGCTGTAGAA<br>CATCTGAGAATGGCTGGGATT |
| RARE -1357<br>RARE -1310<br>RARE -1255<br>RARE -1244 | CTTTCCCAATTGCCAGAAGTG | GATTGCAACATTCTGGAGAGC |
| RARE -1357<br>RARE -1310 | GTTCACTGAGCTGGTCTCATATTC | GCAGGAGTGTGGGAGAGT |
| RARE -1357 | CCCAGCCATTCTCAGATGATATAG | GTGGTGCTGGCACAGAG |
| RARE -1310<br>RARE -1255<br>RARE -1244 | GCTCATGACCCTGGGAATAG | GGTCCCATCTCTCACCAATAG |

### Table S4 (cont.)

| Target Sequence in mSAA3 promoter | Forward Primers (5' to 3') | Reverse Primers (5' to 3') |
|---|---|---|
| RARE -1160<br>RARE -1134<br>RARE -1058 | GAATGTTGCAATCAGTGAGGAG | CAGCCTGAGATGATGGTGAA |
| RARE -1160 | CTGCTGCTATTGGTGAGAGATG | TGTTATCACTAGAGGACATGGAGATA |
| RARE -1058 | GGGATATCTCCATGTCCTCTAGT | GTGGAGCAGCTTGAGCATTA |
| RARE -689<br>RARE -603 | CTTCAAGCTAGGATGAACAGAGG | GGGAAAGAGAGAAGAAAGCATCA |
| RARE -689 | CTGTGCTGCCTGGATATGAT<br>GGTCTGCAGGTGTCTATCTTC | CACCTTGATGTTGGCATTGTT<br>CCCTGGCAATGGGCATA |
| RARE -603 | GGAACAATGCCAACATCAAGG | CACACACTGGATTGGATGGA |
| RARE -463 | GTGTGAGCCAACTGCTCTTA<br>ACCCAGCTTGATGCTTTCT<br>CCATCCAATCCAGTGTGTGT | ATCTTAGCATGGACGGTGTG<br>GGTGAGAAGCTATTGGCATTTG<br>GCAGAGGAAGGGTTGGTTT |
| RARE -327 | CACACCGTCCATGCTAAGAT<br>CAAATGCCAATAGCTTCTCACC<br>AACCAACCCTTCCTCTGCTA | GGTAGAGGTGATGGTTGACTTC<br>CAAGAGGGTGGACTCACAAG<br>GGACTGTGACAGCATTGCATA |
| RARE -224 | GAAGTCAACCATCACCTCTACC<br>CTTGTGAGTCCACCCTCTTG | AAATATCCTCGGACACACCATC<br>CAAGGTTCGAGGGTCCTTT |
| RARE -116 | GATGGTGTGTCCGAGGATATTT<br>AAAGGACCCTCGAACCTTG | GACAGTGAGACAGATGACACAG<br>AAGAATCTGTGCGACAGTGA |
| RARE-11<br>RARE 15 | CTGTGTCATCTGTCTCACTGTC<br>ACTGTCGCACAGATTCTTCTC | AACTAGCATGCTGTCCTCAAA<br>AGGCTCAGTACCATCCAAAC |
| RARE 15 | CTGTGGGTTGGGATCTTGT | CTTTAGAGTGCTCCTCCAGTG |
| RARB RARE Positive Control | CGGGTAGGGTTCACCGAAAGTTCACTCGCA | TGCGAGTGAACTTTCGGTGAACCCTACCCG |
| H3K27Ac Binding Positive Control | TCTCCCTAAACTCTTCCACTCG | ATGTGCCTAAAACGCATCACTACTA |
| H3K27Ac Binding Negative Control | ACAACTGAGGGGAGGAGAGAAG | GCTGCATTTGTTTTCATTCAGT |

**Table S5: qPCR primers for gene expression assays**

| Target gene | Forward Primer (5' to 3') | Reverse Primer (5' to 3') |
|---|---|---|
| mouse *Gapdh* | TGGCAAAGTGGAGATTGTTGCC | AAGATGGTGATGGGCTTCCCG |
| mouse *Saa1* | ATCACCAGATCTGCCCAGGA | CCTTGGAAAGCCTCGTGAAC |
| mouse *Saa2* | ACCAGATCTGCCCAGGAGAC | GCATGGAAGTATTTGTCTCCATCT |
| mouse *Saa3* | GACATGTGGCGAGCCTACTC | TTGGCAAACTGGTCAGCTCT |
| human *GAPDH* | CCTGGTCACCAGGGCTGCTTTTAAC | GTCGTTGAGGGCAATGCCAGCC |
| human *SAA1* | GGCATACAGCCATACCATTC | CCTTTTGGCAGCATCATAGT |
| human *SAA2* | GCTTCCTCTTCACTCTGCTCT | TGCCATATCTCAGCTTCTCTG |

**Table S6. siRNA probe sequences for targeted gene silencing**

| Target gene | Probe Sequences |
|---|---|
| human *RARA* | GCAAAUACACUACGAACAA<br>CCAAGGAGUCUGUGAGAAA<br>GAGCAGCAGUUCUGAAGAG<br>GAACAACGUGUCUCUCUGG |
| human *RARB* | CAGCUGAGUUGGACGAUCU<br>CGAGAUAAGAACUGUGUUA<br>GGCCUUACCCUAAAUCGAA<br>UCACAGAUCUCCGUAGCAU |
| human *RARG* | GAAAGGGCCAUUACUCUGA<br>CAAGGAAGCUGUGCGAAAU<br>GGAGAACCCUGAAAUGUUU<br>UAGAAGAGCUCAUCACCAA |
| mouse *Rara* | GCAAGUACACUACGAACAA<br>AAGACAAAUCAUCCGGCUA<br>CGGUGCGAAACGAUCGAAA<br>CGAAUCUGCACGCGGUACA |
| mouse *Rarb* | GAUAAGAACUGCGUCAUUA<br>GAAAGGUGCCGAACGUGUA<br>GAUCUACACUUGCCAUCGA<br>AAGAGUCUGUUAGGAAUGA |
| mouse *Rarg* | GUAAGGAACGAUCGAAACA<br>GCGGAUCUGUACAAGGUAU<br>GCAGGACACUAUGACAUUC<br>GGAGCAGGCUUCCCAUUCG |
